# Coregulation of tandem duplicate genes slows evolution of subfunctionalization in mammals

**DOI:** 10.1101/019166

**Authors:** Xun Lan, Jonathan K. Pritchard

## Abstract

Gene duplication is a fundamental process in genome evolution. However, most young duplicates are degraded into pseudogenes by loss-of-function mutations, and the factors that allow some duplicate pairs to survive long-term remain controversial. One class of models to explain duplicate retention invokes sub- or neofunctionalization, especially through evolution of gene expression, while other models focus on sharing of gene dosage. While studies of whole genome duplications tend to support dosage sharing, the primary mechanisms in mammals–where duplications are small-scale and thus disrupt dosage balance– are unclear. Using RNA-seq data from 46 human and 26 mouse tissues we find that sub-functionalization of expression evolves slowly, and is rare among duplicates that arose within the placental mammals. A major impediment to subfunctionalization is that tandem duplicates tend to be co-regulated by shared genomic elements, in contrast to the standard assumption of modularity of gene expression. Instead, consistent with the dosage-sharing hypothesis, most young duplicates are down-regulated to match expression of outgroup singleton genes. Our data suggest that dosage sharing of expression is a key factor in the initial survival of mammalian duplicates, followed by slower functional adaptation enabling long-term preservation.

## Main Text

Gene duplications are a major source of new genes, and ultimately of new biological functions (*1–4*). However, new duplicates are usually functionally redundant and thus susceptible to loss-of-function mutations that degrade one of the copies into a pseudogene. The average half-life of new primate duplicates has been estimated at just 4 million years (*5*). This raises the question of what are the evolutionary forces governing the initial spread and persistence of young duplicates.

There has been a great deal of work to understand why some duplicate pairs do survive over long evolutionary timescales (*6*). One class of explanations–the dosage-balance models–focus on the importance of maintaining correct stoichiometric ratios in gene expression between different genes (*4, 7–11*). These models are especially attractive following whole genome duplication (WGD), as WGD should maintain the original dosage ratios of all genes. Subsequent gene losses would thus disrupt dosage balance. Several studies support the importance of dosage sharing following WGD (*4, 12, 13*).

A second class of explanations focuses on functional partitioning of duplicates, either by neofunctionalization (one copy gains new functions) or subfunctionalization (the copies divide the ancestral functions between them). An influential model known as Duplication-Degeneration-Complementation proposes that complementary degeneration of regulatory elements leads to complementary expression of the two copies in different tissues, such that both copies are required to provide the overall expression of the ancestral gene (*14*). Similarly, neofunctionalization of expression could lead to one gene copy gaining function in a tissue where the parent gene was not expressed. Functional divergence may also occur at the protein level (*15*), but this is generally thought to be a slow process, so that divergence usually starts through changes in regulation (*1, 16*).

It is currently unclear which mechanisms are most important for long-term survival of mammalian duplicates. In mammals, most duplications arise through segmental duplications or retrotranspositions that increase copy numbers of just one or a few genes. Hence they are likely to immediately disrupt dosage balance, and to the extent that dosage balance is important, it would seem to favor gene loss rather than preservation. Thus, from a theoretical point of view, DDC remains an attractive explanation in mammals, but has not been fully tested.

We therefore set out to test whether recent gene expression data from many tissues in human and mouse support the model of duplicate preservation by sub- or neofunctionalization of expression. To this end, we analyzed RNA-seq data from ten individuals for each of 46 diverse human tissues collected by the GTEx Project (*17*), and replicated our main conclusions using RNA-seq from 26 diverse mouse tissues (*18*).

We first developed a computational pipeline for identifying duplicate gene pairs in the human genome (Supp. Inf. Section 2). After excluding annotated pseudogenes, we identified 1,444 high-confidence reciprocal best-hit duplicate gene pairs with >80% alignable coding sequence and >50% average sequence identity. These criteria excluded ancient duplicates and most complex multi-gene families. We used synonymous divergence *d_S_* as a proxy for divergence time, while noting that divergence of gene pairs may be downwardly biased due to nonallelic homologous gene conversion in young duplicates (*19*). We estimate that *d_S_* for duplicates that arose at the time of the human-mouse split averages ∼0.4 and that most pairs with *d_S_* >∼0.7 predate the origin of the placental mammals (Figs. S3, S4). Most of our analysis focuses on duplicates that likely arose within the mammals and thus postdate whole genome duplications that occurred in the early vertebrates (*20*).

We next considered GTEx RNA-seq data from 46 tissues. Accurate measurement of expression in gene duplicates can be challenging because RNA-seq reads may map equally well to both gene copies. Alternatively, mapping may be systematically biased if the two copies have differential homology with other genomic locations. To overcome these challenges, we developed a new method specifically for estimating the expression levels of duplicate genes (Supp. Inf. Section 3). In brief, we identified paralogous positions within each duplicate pair for which reads from both copies would map uniquely to the correct gene. Only these positions were used for estimating expression ratios. This approach is analogous to methods for measuring allele-specific expression (*21*). These strict criteria mean that some very young genes are excluded from our expression analyses as unmappable but, for the remaining genes, simulations show that our pipeline yields highly accurate, unbiased estimates of expression ratios (Fig. S1).

**Figure 1.**
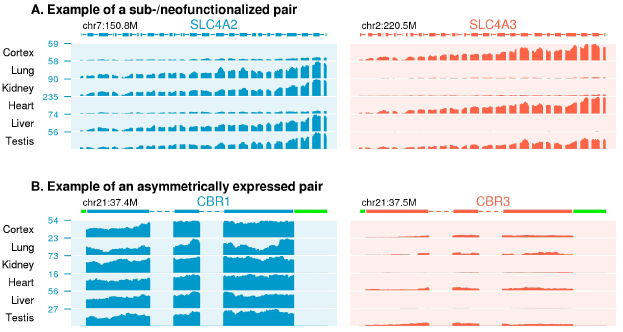
Expression profiles of duplicate genes. **A**. *A gene pair whose expression profile is consistent with sub- or neofunctionalization: i.e., each gene is significantly more highly expressed than the other in at least one tissue*. **B**. *An asymmetrically expressed gene pair. Notice that expression of CBR1 exceeds expression of CBR3 in all tissues. Introns have been shortened for display purposes. The Y-axis shows read depth per billion mapped reads. Green regions in the gene models are unmappable*.

This new read mapping pipeline allowed us to classify gene pairs into categories based on their co-expression patterns (Supp. Inf. Sections 3, 6). We classified a gene pair as being potentially sub-/neofunctionalized if both gene copies are significantly more highly expressed than the other in at least one tissue each (at least 2-fold difference and p<0.001; example in Fig. 1A). We also noticed that many gene pairs show asymmetric patterns of gene expression, in which one gene tends to have higher expression than the other. For all duplicates, we classified the gene with higher overall expression as the “major” gene, and its partner as the “minor” gene. We refer to pairs with consistent asymmetry as *asymmetrically expressed duplicates* (AEDs; example Fig. 1B). Pairs were classified as AEDs if the major gene was significantly more highly expressed in at least 1/3 of tissues where either gene is expressed, and not lower expressed than its partner in any tissue. The remaining duplicates were classified as *no difference* pairs, though many show weaker levels of asymmetry.

Analysis of the RNA-seq data indicates that few duplicate pairs show evidence of sub-/neofunctionalization of expression (Fig. 2A-C). Moreover, most gene pairs with such patterns are very old, dating to before the emergence of the placental mammals: for duplicates with *d_S_* < 0.7, just 15.2% of duplicates are classified as potentially sub-/neofunctionalized in expression. Given that even modest variation in expression profiles across tissues would meet our criteria for subfunctionalization, the fraction of truly subfunctionalized duplicates may be even lower.

In a separate analysis, we found similar levels of apparent subfunctionalization in a mouse dataset (*18*) with better representation of fetal tissues (Fig. S11). We also wondered whether subfunctionalization might instead occur through differential splicing of exons (*22*); however we found little evidence for this (Fig. S13). Lastly, we hypothesized that subfunctionalization might be more prevalent in gene pairs with higher tissue specificity (as they may have more tissue-specific enhancers) but this is not the case (Fig. S10).

While relatively scarce, the genes identified as potentially subfunctionalized do exhibit systematic differences from other duplicates. First, subfunctionalized gene pairs are expected to be under stronger selective constraint than genes without diverged expression, since the two copies are not functionally redundant. Consistent with this, we find that putatively subfunctionalized genes tend to have a higher fraction of rare variants in human polymorphism data (*23*) (p=2×10^−5^ for missense mutations; Fig. 2D). Second, we hypothesized that if subfunctionalized genes have distinct functions, then they should often be associated with distinct genetic diseases. Using a database of gene associations for diverse diseases (*24*) we found a strong correlation between the degree of expression subfunctionalization and the number of diseases reported for one member of the gene pair only (p=5×10^−12^ controlling for relevant covariates; Fig. 2E, Table S3).

**Figure 2.**
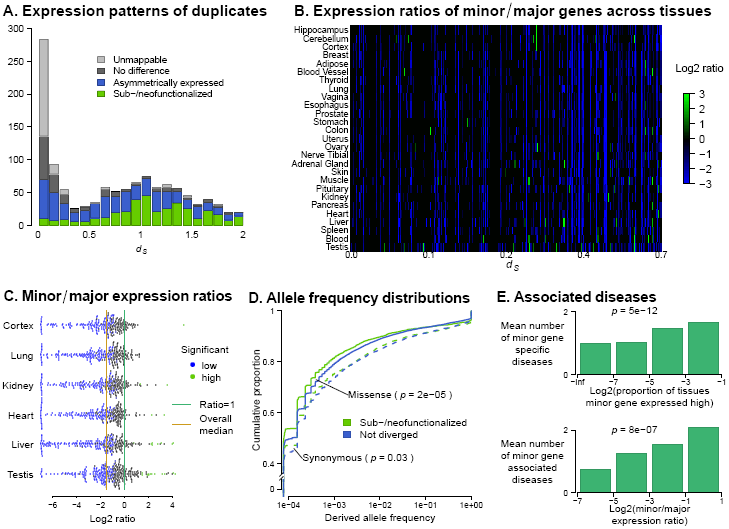
Properties of subfunctionalized genes. **A**. *Classification of gene pairs by expression patterns. For context, note that duplicates arising at the human-mouse split would have ds ∼0.4*. **B**. *Heat map of expression ratios for duplicate pairs. For each duplicate pair (plotted in columns) the ratios show the tissue-specific expression level of the minor gene relative to its duplicate. Green indicates evidence for subfunctionalization; consistently blue columns indicate AEDs. Black indicates tissue ratios not significantly different from 1 (p>.001)*. **C**. *Distributions of expression ratios in different tissues (minor genes/major genes). Ratios significantly >1 marked in green*. **D**. *Selective constraint on subfunctionalized genes. Frequency spectra of human polymorphism data* (*23*) *for synonymous and nonsynonymous variants in subfunctionalized duplicates (green) and duplicates without significant expression differences (black). The plots show cumulative derived allele frequencies at segregating sites. The lines that climb more steeply (subfunctionalized genes) have a higher fraction of rare variants, indicating stronger selective constraint*. **E**. *Disease burden of minor genes is highly correlated with degree of subfunctionalization (top) and overall expression relative to major genes (bottom). Note: data in B, C and D are for ds* < 0.7.

In sharp contrast to the expectations of subfunctionalization, many duplicate pairs instead have systematically biased expression, as seen previously in some systems following whole genome duplication (*25, 26*). Across all duplicate pairs, the mean expression of the less-expressed gene is 40% that of its duplicate (Fig. 2B, C; p∼0 relative to a model with no true asymmetry). Among duplicates that likely arose within the placental mammals (*d_S_* < 0.7), 52.6% of duplicates are AEDs, compared to just 15.2% potentially subfunctionalized pairs. As might be expected, the minor genes at AEDs show clear evidence of evolving under reduced selective constraint relative to their duplicate partners, both within the human population (Fig. S16) and between species (Fig. S14). Furthermore, in gene pairs with asymmetric expression the minor genes tend to be associated with significantly fewer diseases (p=8×10^−7^; Fig. 2E). Nonetheless, despite their reduced importance, minor genes are not dispensable: 97% of minor genes have *d_N_/d_S_*<1, which is a hallmark of protein-coding constraint (Fig. S14).

Together, these results suggest that subfunctionalization of expression evolves relatively slowly. To better understand why this is, we explored which genomic features are correlated with divergent expression profiles of duplicate genes (Fig. 3). Controlling for *d_S_*, the most important predictor of sub-/neofunctionalization is that the duplicates are located on different chromosomes. Most duplicate pairs arise as segmental duplications (*2, 27*) and are found close together in the genome: 87% of young gene pairs (*d_S_* <0.1) are on the same chromosome (Fig. 3A). The duplicates may subsequently become separated as the result of chromosomal rearrangements (*28*), however this is a slow process–it is not until *d_S_*=0.6 that half of gene duplicates are on different chromosomes.

Our data suggest that genomic separation of gene duplicates is a major factor enabling expression divergence. Duplicates that are separated in the genome are much more likely to be sub-/neofunctionalized (p=5×10^−23^, Fig. S17) and tend to have less correlated expression profiles across tissues (Fig. 3B). When the duplicates are on the same chromosome, there is also a significant, though weaker, effect of the distance between the genes and their expression correlation (p=0.002, Fig. S19). In contrast, the asymmetry of mean expression is uncorrelated with whether the duplicates are in cis or trans (p=0.9, controlling for d_S_).

**Figure 3.**
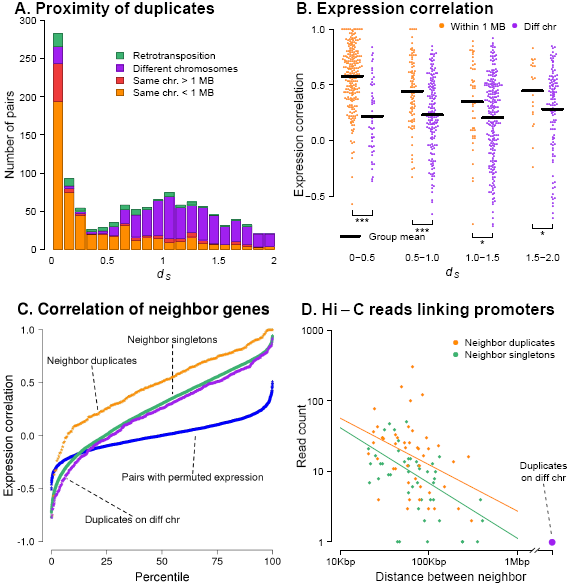
Co-regulation of tandem duplicates. **A**. *Numbers of duplicate pairs in cis and trans, as a function of ds, showing that most young pairs are nearby in the genome*. **B**. *Correlation of expression profiles of duplicates across tissues, for tandem and separated pairs*. **C**. *Overall distributions of correlations for different classes of genes*. **D**. *Numbers of Hi-C links between neighboring gene pairs. (Gene pairs within 20kb excluded due to limited resolution of the assay; singleton pairs randomly downsampled for plotting.)*

These results echo previous observations that, in general, genes that are close in the genome tend to have correlated expression (*29*) and frequently share eQTLs (*30*). However, this effect is especially strong for duplicates: gene expression is more correlated for tandem duplicates than for singleton neighbors (p=1×10^−19^, Fig. 2C) and duplicates share eQTLs at higher rates than matched singletons in two data sets (p=6×10^−4^ and 5×10^−4^, Supp. Inf. Section 12) (*30, 31*).

We hypothesized that since promoters of duplicate genes share related sequences, they may often share the same functional elements. We thus used Hi-C data to explore the regulatory links among duplicates (*32*). Current Hi-C data have limited resolution for assessing enhancer-promoter connectivity; nonetheless we observed a weak signal that individual enhancers are more frequently linked to both promoters of duplicate pairs than to both promoters of matched singleton pairs (p=0.02, Supp. Inf. Section 12). Further, we found a strong signal of direct promoter-promoter links between duplicate genes (Fig. 2D). Pairs of both nearby singleton and duplicate genes frequently show high numbers of promoter-promoter read pairs, however duplicate pairs have systematically higher numbers of links (p=3×10^−6^, Supp. Inf. Section 12). Trans-duplicates show no evidence for Hi-C linkages. The large numbers of promoter-promoter links may reflect a tendency of co-regulated genes to be transcribed simultaneously within transcription factories (*33*).

Thus far, our results argue that expression subfunctionalization evolves slowly, in large part because tandem duplicates tend to be co-regulated. As noted above, an alternative explanation for the initial survival of duplicates is that they are both necessary to produce the required expression dosage (*4*). However, in contrast to whole genome duplications, the small-scale duplications that are typical in mammals would initially disrupt dosage of the duplicated genes relative to all other genes. Thus, if dosage sharing is important in mammals, this would suggest that following tandem duplication, the duplicates should rapidly evolve reduced expression. Subsequent loss of either gene would cause a deficit of expression and thus be deleterious.

**Figure 4.**
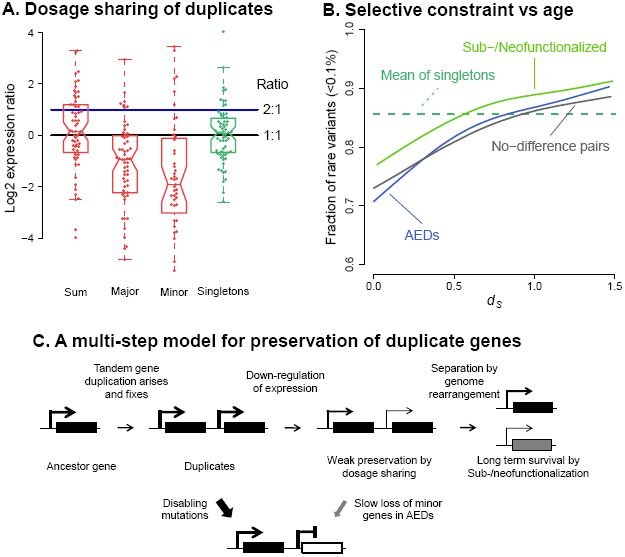
Long-term survival of duplicate genes. **A**. *Rapid evolution of dosage sharing. Expression levels of young duplicates compared to their macaque orthologs in 6 tissues (*34*), for human duplicates that are single-copy genes in macaque. Sum shows the summed expression of both duplicates, relative to expression of the macaque orthologs in the same tissues. “Major” and “Minor” show corresponding ratios for major and minor genes separately, classified using GTEx data. The green data show a random set of singleton orthologs. Each tissue-gene expression ratio is plotted separately*. **B**. *The strength of purifying selection in humans increases with duplicate age. The fraction of rare missense variants in a large human data set* (*23*) *is used as a proxy for the strength of purifying selection*. **C**. *Conceptual model of duplicate gene evolution. Other transitions not explicitly shown would occur at lower but nonzero rates*.

To evaluate this, we analyzed the expression of human duplicates that arose since the human-macaque split, using RNA-seq data from 6 tissues in human and macaque (*34*) (Fig. 4A, Supp. Inf. Section 7). Indeed, there is a very clear signal that both human copies tend to evolve reduced expression, such that the median summed expression of the human duplicates is close to the expression of the singleton orthologs in macaque (median expression ratio 1.11; this is significantly less than the 2:1 expression ratio expected based on copy number, p=8×10^−6^). Thus, dosage sharing may be a frequent first step in the preservation of tandem duplicates. However, although dosage sharing evolves quickly, it is notable that duplicate genes tend to remain relatively unconserved over long evolutionary timescales (*d_S_* ≤ 0.7 or roughly the age of placental mammals; Fig. 4B, S15).

In summary, we have reported here that sub-/neofunctionalization of expression occurs slowly for most gene pairs, and generally does not happen until the duplicates are separated by genomic rearrangements. These observations imply that the tissue expression profiles of tandem duplicates are not fully modular, as often assumed. In contrast however, we observe widespread differences in mean expression levels between duplicates, and the degree of asymmetry does not depend on whether the duplicates are in cis or trans. This suggests that mean expression may be more free to evolve independently through changes in promoter strength.

Figure 4C summarizes a conceptual multi-step model for the evolution of mammalian duplicates. We propose that downregulation is often a key first step enabling the initial survival of duplicates, followed by dosage sharing as suggested for WGDs (*4*). Subsequently, the relative expression levels of the two genes evolve as a random walk, but do so slowly due to constraint on their combined expression (*13, 35*). If expression becomes asymmetric, this reduces functional constraint on the minor gene and may lead to gene loss. We propose that genomic separation is often a second key step in long-term survival, as this frees the expression of the duplicates to evolve independently, and may also encourage protein adaptation (*36*). These additional steps enable true functional differentiation and long-term survival of duplicates.

## Acknowledgments

This work was funded by NIH grants ES025009 and MH101825, and by the Howard Hughes Medical Institute. We thank Hunter Fraser for prepublication access to data, and Hunter Fraser, Audrey Fu, Arbel Harpak, Yang I. Li, Dmitri Petrov, Patrick Phillips, Molly Przeworski, and Arlin Stoltzfus for comments and discussion.

